# Improving Structural Plausibility in 3D Molecule Generation via Property-Conditioned Training with Distorted Molecules

**DOI:** 10.1101/2024.09.17.613136

**Authors:** Lucy Vost, Vijil Chenthamarakshan, Payel Das, Charlotte M. Deane

**Affiliations:** Department of Statistics, University of Oxford, Oxford, UK; IBM Research, Yorktown Heights, New York, USA

## Abstract

Traditional drug design methods are costly and time-consuming due to their reliance on trial-and-error processes. As a result, computational methods, including diffusion models, designed for molecule generation tasks have gained significant traction. Despite their potential, they have faced criticism for producing physically implausible outputs. We alleviate this problem by conditionally training a diffusion model capable of generating molecules of varying and controllable levels of structural plausibility. This is achieved by adding distorted molecules to training datasets, and then annotating each molecule with a label representing the extent of its distortion, and hence its quality. By training the model to distinguish between favourable and unfavourable molecular conformations alongside the standard molecule generation training process, we can selectively sample molecules from the high-quality region of learned space, resulting in improvements in the validity of generated molecules. In addition to the standard two datasets used by molecule generation methods (QM9 and GEOM), we also test our method on a druglike dataset derived from ZINC. We use our conditional method with EDM, the first E(3) equivariant diffusion model for molecule generation, as well as two further models—a more recent diffusion model and a flow matching model—which were built off EDM. We demonstrate improvements in validity as assessed by RD-Kit parsability and the PoseBusters test suite; more broadly, though, our findings highlight the effectiveness of conditioning methods on low-quality data to improve the sampling of high-quality data.

## 1 Introduction

Drug design involves complex optimisation steps to obtain molecules that achieve desired biological responses. Traditional methods rely on trial-and-error, leading to high costs and limited productivity [25]. Computational approaches, especially deep learning models, aim to reduce costs and expedite processes by reducing failures. One way that such models aim to do this is by generating molecules with desirable properties, particularly in terms of binding to their target. To achieve this, a model must first master the fundamental task of generating structurally viable molecules.

While many models historically operated in 1D or 2D space [20, 5, 6], focus has recently shifted towards developing models capable of directly outputting both atom types and coordinates in 3D. Autoregressive models were once prominent in this domain, generating 3D molecules by adding atoms and bonds iteratively [14, 22, 16]. However, such models suffer from an accumulation of errors during the generation process and do not fully capture the complexities of real-world scenarios due to their sequential nature, potentially losing global context [12, 13]. To address these limitations, recent studies have turned to diffusion models, which iteratively denoise data points sampled from a prior distribution to generate samples. Unlike autoregressive models, diffusion-based methods can simultaneously model local and global interactions between atoms. Nevertheless, diffusion in molecule generation has faced criticism for yielding implausible outputs [8, 3]. There have been ongoing efforts to improve the performance of models trained on small molecules such as those found in the QM9 dataset [19, 10, 15, 24, 11], but achieving success in generating larger molecules, as encountered in datasets like GEOM [1], remains challenging without incorporating additional techniques such as energy minimisation or docking [27].

In this paper, we focus on enhancing the ability of a diffusion model to generate plausible 3D druglike molecules. To achieve this, we use the property-conditioning method developed by Hoogeboom *et al*. [10]. Instead of conditioning a model on pre-existing properties, we condition on conformer quality, training the model to not only generate molecules, but also to distinguish high- and low-quality chemical structures.

To achieve this, we generate distorted versions of each of the three datasets we evaluate the method on: QM9, GEOM, and a subset of ZINC. We sample molecules from each dataset and apply random offsets to their original coordinates, based on a maximum distortion value. Each distorted molecule is assigned a label representing the degree of warping applied and is added back to the dataset. Non-distorted molecules are also labeled, identifying them as high-quality conformers. Using these datasets of molecules with varying levels of quality, we train property-conditioned models, encouraging the model to learn to label molecule validity while simultaneously training it to generate molecules.

First, we evaluate our conditioning method with EDM, the first E(3) equivariant diffusion model for molecule generation [10]. We then test it on two additional models: a geometry-complete diffusion model [15] and a flow matching method [21], both designed to enhance the structural plausibility of generated molecules. We also employ two datasets of druglike molecules: the GEOM dataset, and another derived from the ZINC database. This ensures the method extends beyond the scope of smaller molecules found in QM9.

Our findings demonstrate that across the models tested, conditioning a model with low-quality conformers enables it to discern between favourable and unfavourable molecular conformations. This allows us to target the area of the learned space corresponding to high-quality molecules, resulting in an improvement of the validity of generated molecules. More broadly, this demonstrates the potential of supplementing molecule generation methodologies not solely with examples of desired molecules but also with instances exemplifying undesired outcomes.

## 2 Methods

### 2.1 Generation of 3D molecules

Hoogeboom *et al*.[10] introduced the first E(3)-equivariant diffusion model (EDM) for generating 3D small molecules. Since then, significant efforts have been made to modify the original EDM, whether to adapt the method for structure-based drug design [4, 7, 12] or to enhance the validity of the generated molecules [17]. Notable examples of the latter include GCDM (Geometry-Complete Diffusion Model) [15] and EquiFM [21]. GCDM addresses the limitations of diffusion models that rely on molecule-agnostic and non-geometric graph neural networks (GNNs) for 3D graph denoising by introducing a geometry-complete approach. In contrast, EquiFM focuses on the issue of unstable probability dynamics in existing diffusion models by incorporating geometric flow matching, merging the advantages of equivariant modeling with stabilised probability dynamics.

### 2.2 Conditioning on conformer quality

The authors of EDM developed an extension to their method to carry out conditional molecule generation. In this instance, property annotations are included alongside each of the molecules in the training dataset, and at inference, molecules can be generated with a desired value of this property. We use this property-conditioning method to train models conditioned on conformer quality. To implement this, we first generated datasets with 3D conformers of molecules of variable quality levels, and corresponding annotations. We generated distorted versions of a subset of molecules from each of the datasets we used. For each molecule, its 3D coordinates, represented as *C* = {(*x*_*i*_, *y*_*i*_, *z*_*i*_)} where *i* denotes the atom index, were obtained. Subsequently, a random number *D* within the range of 0 to *D*_*max*_ angstroms, labelled as the maximum distortion, was sampled:

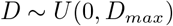

This value represents the maximum distance in angstrom that could be added to atoms in that molecule: in other words, the sampled distortion value determines the maximum extent of perturbation to be applied to the molecule’s structure. Following this, random offsets were generated within the range of 0 to the sampled distortion, *D*, for each dimension of every atom’s coordinates:

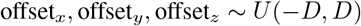

These offsets were then applied to the original coordinates:

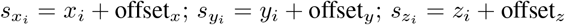

Resulting in a ‘distorted’ version of the molecule. This distorted molecule, along with its corresponding sampled distortion value *D*, was subsequently added to the training set. Following the generation of the distorted datasets, we use the property-conditioning training protocol outlined by Hoogeboom *et al*. to train on them, using the distortion factor *D* as the property of interest, and follow the sampling protocol to generate molecules corresponding to *D* = 0Å.

We assessed all models by their ability to generate molecules that pass RDKit sanitisation and the PoseBusters suite (see SI).

## 3 Results and discussion

In this section, we evaluate the performance of EDM, both with and without conditioning, on QM9, GEOM_no h_ and ZINC (see SI for full descriptions). We begin by training the non-conditioned model on all datasets and evaluating them as outlined above. Subsequently, we train conditioned models for all three datasets: QM9, GEOM_no h_, and our ZINC subset. Finally, we assess the broader applicability of the quality conditioning method by assessing it with GCDM and EquiFM.

### 3.1 Performance with no conditioning

First, we assess the performance of EDM when trained without conditioning on all three datasets. For QM9, we use the pretrained model provided by Hoogeboom *et al*. Table 1 shows that the QM9 dataset—comprised of smaller-than-druglike molecules—has the highest baseline model performance, with RDKit and PoseBusters pass rates of 91% and 81%, respectively (see SI for a full breakdown of the results). The baseline model also has a high performance when trained on the GEOM_no h_ dataset, with pass rates of 90% and 70%.

**Table 1:** Performance comparison of EDM trained on diverse molecular datasets using the baseline model with no conditioning, and EDM conditionally trained on distortion factor using a dataset generated with *D*_*max*_ = 0.25Å and 1:50 distorted molecules, and sampled with *D*=0Å. The highest performance for each dataset is shown in bold.

| Dataset |  | Pass Rate |  |
| --- | --- | --- | --- |
|  |  | RDKit | PoseBusters |
| QM9 | baseline | **91%** | **81%** |
|  | conditioned | 73% | 53% |
| GEOM<sub>no h</sub> | baseline | 90% | 70% |
|  | conditioned | **97%** | **81%** |
| ZINC | baseline | 64% | 40% |
|  | conditioned | **90%** | **63%** |

The non-conditioned model trained on the ZINC subset exhibits a much lower RDKit sanitisation pass rate of 64%, and relatively poor PoseBusters pass rate of 40%. Unlike EDM trained on GEOM_no h_, which mostly faced connectivity issues, the most common failures of the molecules from the ZINC subset trained model are in bond lengths and bond angles, passing tests for only 44% and 48% of occurrences, respectively (see SI for a full breakdown of the results).

This decrease in performance compared to GEOM_no h_ could be due to the increased diversity of the dataset: our ZINC dataset does not include repeat conformers of the same molecule, whereas GEOM_no h_ includes up to 30 conformers of each unique molecule. The model might therefore allocate more attention to learning atom types—designing the actual molecule—as opposed to improving its capacity for 3D conformer generation. However, a potentially more likely explanation is the actual molecules that make up the GEOM_no h_ dataset. The ZINC subset comprises entirely medium-sized compounds, whereas GEOM_no h_, in addition to containing some medium-sized molecules, also incorporates the entirety of QM9, and as Table 1 shows, EDM has a high performance using the QM9 dataset. The inclusion of QM9 in GEOM_no h_ also means that the average molecule size is smaller than that of ZINC subset, which may also give EDM an advantage on GEOM_no h_ type molecules compared to the larger molecules in ZINC.

Overall, our findings indicate that while the baseline, non-conditional EDM model demonstrates proficiency in generating small to medium-sized compounds, its performance deteriorates on a dataset comprising of exclusively medium-sized molecules, resulting in the generation of numerous molecules with physically implausible bond lengths and angles. We next explore steering the model away from generating physically implausible molecules via our conditioning method.

### 3.2 Conditioning on distortion factor

To determine a sensible proportion of distorted molecules and the degree of distortion for effective conditional training, we conducted ablation studies using the GEOM_no h_ dataset. Of the parameters tested, the best performance was observed using a ratio of 1:50 distorted molecules and a maximum distortion of 0.25Å (see SI for all ablation test results). Building upon this insight, we generated distorted versions of the remaining datasets, QM9 and ZINC, with these parameters. We then trained conditioned models on each of these modified datasets, sampled 100 molecules from each, and evaluated their quality using RDKit and PoseBusters (Table 1).

When using the QM9 dataset, the highest-performing molecules on both RDKit and PoseBusters tests were generated using the non-conditioned EDM model. Conditional training resulted in a relatively uniform decrease in performance across all PoseBusters tests. This may be attributed to the fact that EDM was specifically developed to perform effectively with QM9. Further, the molecules in this dataset are small (smaller than 9 heavy atoms), which our earlier results suggest facilitates the model’s ability to distinguish between high-quality and low-quality conformers without needing examples of the latter.

The models trained conditionally on GEOM_no h_ and ZINC both generated molecules with improved RDKit sanitisation rates and PoseBusters scores. For GEOM_no h_, as discussed in the ablation tests (see SI), the biggest issues in the baseline were bond angles (80%), followed by bond lengths and internal energy (86%). Conditioning improved performance across all these metrics, reaching 94%, 95%, and 90%, respectively.

Training conditionally with ZINC also resulted in significant improvements across the PoseBusters tests: the lowest pass rates for both the baseline and conditioned models were in bond length/angles, steric clash, and internal energy. The most notable increase in performance for the conditioned model was the improved pass rate for atom connectivity, with this score jumping from 59% to 96%.

Having demonstrated that the conditioning method enhances the structural plausibility of generated molecules when EDM is trained on ZINC or GEOM_no h_, we extend our investigation to determine whether this improvement holds for other models.

### 3.3 Testing the Conditioning Method on Additional Models

To evaluate the broader applicability of our method, we apply it to two other models: GCDM [15] and EquiFM [21]. The performance of these models when trained on GEOM_no h_ and ZINC is presented in Tables 2a and 2b, respectively.

**Table 2:** Performance comparison of GCDM and EquiFM when trained on GEOM_no h_ and our ZINC subset using the default setup (baseline) or conditionally trained on distortion factor using a dataset generated with *D*_*max*_ = 0.25Å and 1:50 distorted molecules, and sampled with *D*=0Å. The highest performance for each dataset is shown in bold.

|  |  | RDKit | PoseBusters |  |  | RDKit | PoseBusters |
| --- | --- | --- | --- | --- | --- | --- | --- |
| GEOM<sub>no h</sub> | baseline | **100%** | 83% | GEOM<sub>no h</sub> | baseline | **97%** | **57%** |
|  | conditioned | 94% | **89%** |  | conditioned | 95% | 44% |
| ZINC | baseline | 78% | 64% | ZINC | baseline | 76% | 39% |
|  | conditioned | **100%** | **78%** |  | conditioned | **96%** | **87%** |

(a) Performance of GCDM
(b) Performance of EquiFM

For both datasets, the molecules generated by GCDM trained with conditioning outperform those produced by the baseline model as assessed with PoseBusters. The improvement observed with the conditioned model is even more pronounced than with the original EDM. This aligns with findings from the GCDM paper, which found that their model generates not only more stable molecules but also more property-specific ones when property conditioning is applied. This suggests that GCDM is more effective at distinguishing between different property values, and in this case, between high- and low-quality regions of the learned space. Training EquiFM using the conditional method does not improve the plausibility of generated molecules when using GEOM_no h_, in which many molecules suffer from connectivity issues (see SI for full breakdown of results). It does, however, improve the plausability of generated molecules when using the ZINC dataset, by a margin similar to that shown by EDM.

In conclusion, our conditioning method that was developed and tested with EDM is able to, without modification, enhance molecular plausibility across different models when looking at GEOM_no h_ and ZINC, with GCDM showing particularly notable improvements. These results suggest that the conditioning approach is broadly applicable and beneficial.

## 4 Conclusions

In this work, we have demonstrated the effectiveness of including low-quality conformers in a training set and conditioning a diffusion model on a label representing conformer quality to enhance the generation of high-quality druglike molecules. By leveraging datasets derived from GEOM and ZINC, alongside a conditioning method proposed by Hoogeboom *et al*., we have successfully improved the validity of generated molecules. Our approach, which focuses on sampling molecules with labels corresponding to low distortion factors, leads to enhancements in RDKit parsability and validity as assessed by PoseBusters for the original EDM, as well as for a subsequent diffusion model, GCDM, and a flow-matching model, EquiFM.

Our findings underscore the importance of considering the quality of conformers in molecule generation processes. The results show that by training models to discern between favorable and unfavorable molecular conformations, we can selectively sample from the high-quality region of learned space, resulting in significant improvements in the validity of generated molecules.

Moving forward, further research could explore additional conditioning methods and datasets to continue improving the quality and diversity of generated molecules. Additionally, investigating the applicability of our approach to other areas of molecular design and exploration could yield valuable insights for drug discovery and beyond. Overall, our study provides a promising avenue for generating valid drug-sized molecules efficiently and effectively.

