## supplementary information for "Improving Structural Plausibility in 3D Molecule Generation via Property-Conditioned Training with Distorted Molecules"

### A Supplementary Information

#### A.1 Method overview

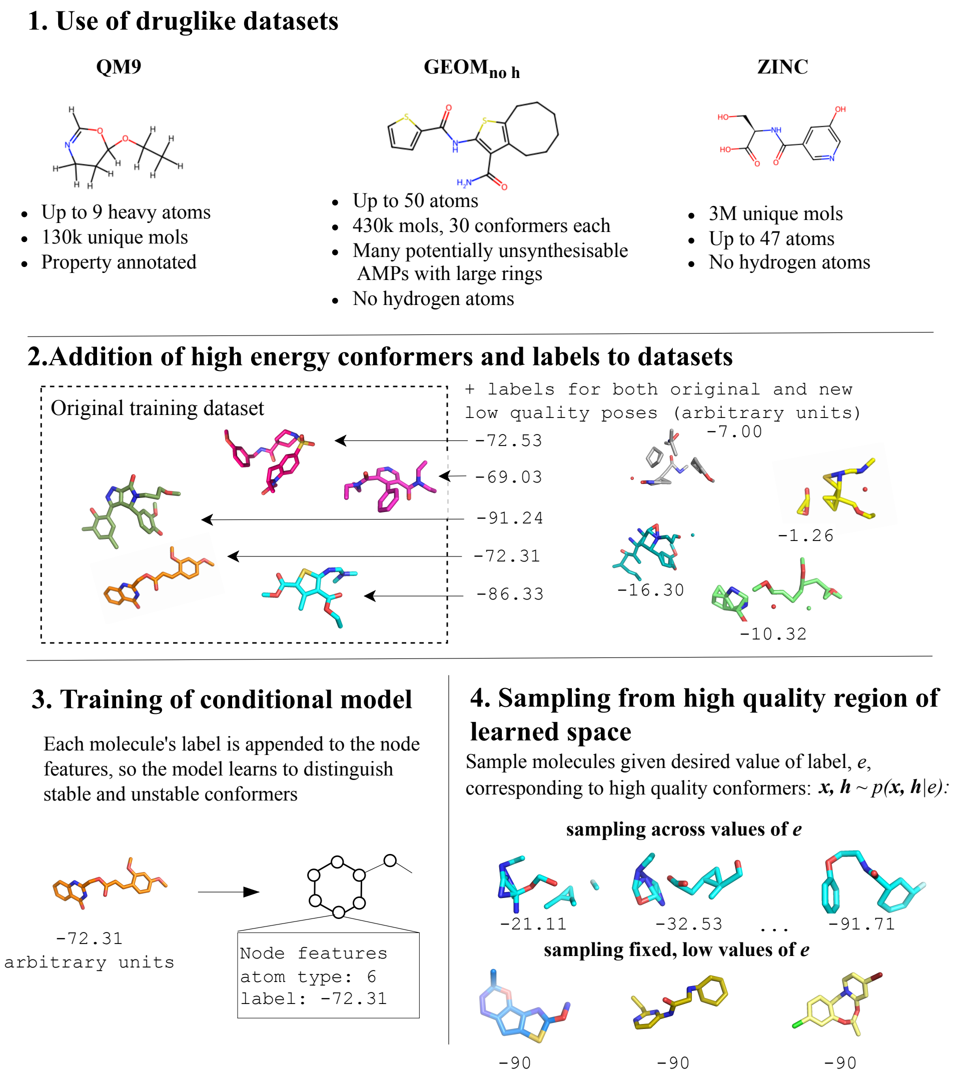


Figure 1: An overall schema of the methods used with (a) the datasets used to train both the unconditional and conditional models, (b) the generation of high energy conformers and their addition to the datasets, (c) the training of the conditional model and (d) the conditional inference

#### A.2 Assessment metrics

For each trained model, we generated 100 molecules. The generated molecules were passed through RDKit, resulting in an RDKit sanitisation pass rate. All molecules were then passed through PoseBusters; however, the first step of the PoseBusters pipeline is to sanitise all molecules, so molecules that fail this automatically fail all subsequent tests. We report the number of molecules that pass RDKit sanitisation as well as all 7 non RDKit sanitisation PoseBusters tests. Finally, we also calculate an internal diversity score with MOSES[18].

##### A.3 Datasets

#### A.3.1 QM9

The QM9 dataset [19] is a widely used benchmark dataset in quantum chemistry and machine learning research. It consists of quantum-mechanical properties and 3D conformers of 130,000 small organic molecules with an average of 17.5 atoms (8.2 heavy atoms).

The QM9 dataset has been extensively used to develop and validate machine learning models for molecular property prediction. However, it has also recently become the central benchmark for *de novo* molecule generation, particularly in the development of diffusion models [10, 15].

###### A.3.2 GEOM

While QM9 features only smaller-than-druglike molecules, GEOM [1] is a larger-scale dataset of molecular conformers. It features 430,000 molecules, of which 317,928 are mid-sized organic molecules from AICures and MoleculeNet [26], and 133,258 molecules are from QM9, resulting in an average molecule size of 44.4 atoms (20.1 heavy atoms). For each molecule, a variable number of conformers are given along with their approximate internal energy as calculated with XTB [2]. From this dataset, Hoogeboom *et al.* [10] retain the 30 lowest energy conformations for each molecule in their work.

Similar to Peng et al. [17], we use a version of GEOM drugs from which hydrogen have been removed ($\text{GEOM}_{\text{no h}}$), as the positions of hydrogen atoms can often be inferred with a high level of confidence [9]. This not only reduces the computational demand of training, but also facilitates more effective learning of heavy atom placements. This leads to the $\text{GEOM}_{\text{no h}}$ dataset becoming the quickest to train on among the three druglike datasets. We therefore use the $\text{GEOM}_{\text{no h}}$ dataset for conducting ablation tests.

###### A.3.3 ZINC

ZINC [23] is a database of commercially-available compounds containing over 230 million purchasable compounds in ready-to-dock, 3D formats.

We generate a training set by selecting a subset of 660,000 molecules from the druglike catalog of the ZINC database. Unlike GEOM, this subset is curated without repeat conformers. Hydrogen atoms are not included, and the average molecule comprises 26.8 heavy atoms.

##### A.4 Ablation tests

To identify the optimal proportion of distorted molecules and the required degree of distortion for effective conditional training, we performed ablation studies using the $\text{GEOM}_{\text{no h}}$ dataset. This dataset was selected due to its inclusion of drug-like molecule sizes, unlike QM9, whilst being a more computationally tractable set to train than the ZINC dataset.

We introduced varying numbers of distorted molecules at different distortion levels (ranging from 0Å, indicating no distortion, to the maximum distortion, $D_{max}$Å) into the original $\text{GEOM}_{\text{no h}}$ dataset. We defined dataset ratios based on the number of distorted and original molecules: for example, a 1:50 ratio indicates one distorted molecule was added for every fifty original molecules. We evaluated each model's performance by training conditioned models and sampling 100 molecules, ensuring that the samples were from the low-distortion-factor region of the learned space (formally, enforcing $D = 0$Å).

The model trained on a dataset with a ratio of 1:50 distorted molecules and a maximum distortion of 0.25Å exhibited the joint highest RDKit parsability rate of 97%, and the highest PoseBusters pass rate at 81%. While several models reached 97% RDKit sanitisation rates (namely 1:20, $D_{max} = 0.5$Å and 1:50, $D_{max} = 0.5$Å), these models exhibited slightly lower PoseBusters pass rates (75% and 78%, respectively). Increasing or decreasing $D_{max}$ further resulted in PoseBusters performance decreasing across all ratios, primarily due to failures in the internal energy test.

This observation suggests that if the training includes molecules that are too distorted, the model does not effectively learn to distinguish between subtly flawed and acceptable molecular structures.

| Maximum distortion permitted, $D_{max}$ (Å) | Ratio of distorted:non-distorted molecules | | | | | |
|---|---|---|---|---|---|---|
| | 1:20 | | 1:50 | | 1:100 | |
| | Pass Rate | | Pass Rate | | Pass Rate | |
| | RDKit | PoseBusters | RDKit | PoseBusters | RDKit | PoseBusters |
| 0.1 | 96% | 73% | 96% | 77% | 96% | 77% |
| 0.25 | 95% | 52% | **97%** | **81%** | 96% | 77% |
| 0.5 | 97% | 75% | 97% | 78% | 95% | 68% |
| 1 | 93% | 57% | 89% | 54% | 62% | 8% |

Table 3: Performance comparison of EDM trained conditionally on $\text{GEOM}_{\text{no h}}$ using a distortion factor, $D$, and sampled with $D$=0Å across various ratios of distorted:non-distorted molecules and maximum distortion values in angstrom.

| Maximum distortion permitted, $D_{max}$ (Å) | Ratio of distorted:non-distorted molecules | | | | | |
|---|---|---|---|---|---|---|
| | 1:20 | | 1:50 | | 1:100 | |
| | Pass Rate | | Pass Rate | | Pass Rate | |
| | RDKit | PoseBusters | RDKit | PoseBusters | RDKit | PoseBusters |
| 0.1 | 91% | 46% | **96%** | **53%** | 81% | 26% |
| 0.25 | 81% | 2% | 81% | 2% | 72% | 4% |
| 0.5 | 49% | 0% | 28% | 0% | 38% | 0% |
| 1 | 41% | 0% | 46% | 0% | 29% | 0% |

Table 4: Performance comparison of EDM trained conditionally on $\text{GEOM}_{\text{no h}}$ using a distortion factor, $D$, and sampled with $D$=$D_{max}$Å across various ratios of distorted:non-distorted molecules and maximum distortion values in angstrom.

Distorted molecules should therefore still bear some resemblance to realistic conformers, albeit with deliberately infeasible bond lengths and angles. On the other hand, insufficient distortion compromises the effectiveness of the conditioning classifier, and the models struggle to distinguish between high-quality and low-quality conformations, leading to poor performance in generating desirable molecules.

These results demonstrate the concept of conditioned training on negative data, and give an idea of the extent of distortion and frequency of distorted molecules to add. We used a ratio of 1:50, and $D_{max} = 0.25$Å for all subsequent tests, but note that any dataset would likely benefit from different exact values of these parameters.

We also examined the quality of molecules generated when sampling from the low-quality region of the learned space (formally, $D = D_{max}$Å). The molecules sampled using $D$= $D_{max}$Å are, as expected, worse than both the conditioned models and the baseline model in terms of PoseBusters pass rates, with the highest reaching only 53%. This poor performance is mainly attributed to failures in the internal energy test.

The RDKit parsability rates of certain models' molecules (specifically 1:20, $D = 0.1$Å and 1:50, $D = 0.1$Å) surpass the baseline model. This observation underscores the importance of incorporating comprehensive evaluations, such as those encompassed by the PoseBusters suite, in the assessment of generative models.

Having established the parameters to use for distorted molecules—both in terms of quality and extent of distortion—that should be included in a dataset to conditionally train EDM, and shown that we can conditionally sample from the high and low-quality areas of the learned space, we move on to applying this method to other datasets.

##### A.5 Conditioning on internal energy

The eXtended Tight Binding (XTB) program [2] is a computational chemistry software package used for molecular modeling and simulations. It is based on the semi-empirical tight-binding approach, which approximates the electronic structure of molecules using a simplified set of parameters derived from quantum mechanics. XTB extends traditional tight-binding methods by incorporating additional empirical corrections to improve the accuracy of calculated properties. It is well-suited for studying large molecular systems where the computational cost of more accurate methods such as density functional theory (DFT) becomes prohibitive.

We use the default implementation of XTB to perform singlepoint energy calculations for each of the molecules, both distorted and original, for QM9, $GEOM_{no\ h}$, and our ZINC dataset. For the original GEOM dataset, the original conformers have energy annotations calculated with the same method, so we only carry out this calculation for the distorted molecules. We carry out this process using two different annotation types: a 'distortion factor', a quantity that represents the extent to which the coordinates have been altered, and an internal energy value, obtained by scoring both original and distorted conformers with the extended tight binding program (XTB) [2].

We've observed that for medium-sized, drug-like compounds, conditioning on a distance-based distortion factor results in improvements for RDKit sanitisation and PoseBusters tests. To investigate whether using a potentially more meaningful label—specifically, an internal energy value obtained using XTB—improves the conditioned models, we followed the previous distortion process. For one in every fifty molecules in each dataset, we generated a distorted version by distorting each atom's coordinates by up to 0.25 Å and added these distorted molecules back to the dataset. These distorted molecules were then passed through an XTB single-point energy calculation. The same calculation was applied to all high-quality, non-distorted molecules in all datasets, excluding the original GEOM dataset, which already has energy annotations calculated with XTB. We then trained each of the models conditionally, and once trained, we sampled 100 molecules from each, this time enforcing $D = E_{min}$, where $E_{min}$ is a fixed value for each dataset corresponding to the lowest internal energy annotation of any molecule in it. Then we once again assessed all 100 molecules using RDKit and PoseBusters (Table 5).

As observed when conditioning on distortion factor, both QM9 and the original GEOM dataset generated lower quality molecules when conditioning with internal energy was carried out. QM9 saw a further decrease in performance when using internal energy, while molecules generated by the model trained on GEOM saw a slight boost in RDKit performance but still exhibited a PoseBusters pass rate of 0%, ultimately being outperformed by the baseline molecules.

Conversely, conditionally training EDM on $GEOM_{no\ h}$ with internal energy resulted in increased performance compared to both the baseline and the conditioned model using the distance distortion factor. The RDKit and PoseBusters pass rates reached 98% and 84%, respectively. The model trained on ZINC, however, exhibited a decrease in both RDKit and PoseBusters pass rate. These pass rates were also lower than those of the molecules generated with the ZINC model trained conditionally on distortion factor.

When using the distance based distortion factor, we enforced $D$=0Å when sampling, as this represented molecules that had not undergone any distortion. The distortion factor, ranging from 0Å to $D_{max}$Å, provides a straightforward measure of how much the structure of a molecule has been altered. In contrast, internal energy values, although physically meaningful, are challenging to compare directly between different molecules. Each molecule can have a low internal energy corresponding to a high-quality conformer and a high one corresponding to a low-quality conformer, but the lowest energy conformers of some molecules may still have higher energy values than the highest energy annotations of others. This inconsistency likely contributed to the observed performance decreases when conditioning on internal energy.

##### A.6 Full PoseBusters outputs

Below we provide the full outputs for each model's molecules when assessed with PoseBusters.

| Dataset | | RDKit | PoseBusters |
|---|---|---|---|
| QM9 | baseline | **91%** | **81%** |
| | XTB | 57% | 19% |
| GEOM | baseline | **82%** | **24%** |
| | XTB | 34% | 0% |
| $\text{GEOM}_{\text{no h}}$ | baseline | 90% | 70% |
| | XTB | **98%** | **84%** |
| ZINC | baseline | 64% | **40%** |
| | XTB | **65%** | 36% |

Table 5: Performance comparison of EDM trained on diverse molecular datasets using a baseline (the unconditional EDM), and using a conditioned model, for which the model is trained on an XTB internal energy estimate. The highest performance for each dataset is shown in bold.

| Dataset | Sanitisation | All Atoms Connected | Bond Lengths | Bond Angles | Internal Steric Clash | Aromatic Ring Flatness | Double Bond Flatness | Internal Energy | All Tests Passed | Diversity |
|---|---|---|---|---|---|---|---|---|---|---|
| Performance with no conditioning | | | | | | | | | | |
| QM9 | 91% | 100% | 91% | 91% | 91% | 91% | 91% | 81% | 81% | 0.88 |
| GEOM | 82% | 44% | 76% | 74% | 75% | 82% | 82% | 66% | 24% | 0.85 |
| $GEOM_{no\ h}$ | 90% | 90% | 86% | 80% | 91% | 90% | 89% | 86% | 70% | 0.85 |
| ZINC | 64% | 59% | 44% | 48% | 49% | 64% | 64% | 51% | 40% | 0.84 |
| Ablation tests | | | | | | | | | | |
| 1:20, $D_{max}$ = 0.1Å | 96% | 93% | 94% | 92% | 96% | 96% | 96% | 82% | 73% | 0.85 |
| 1:20, $D_{max}$ = 0.25Å | 95% | 73% | 88% | 90% | 95% | 95% | 95% | 82% | 52% | 0.85 |
| 1:20, $D_{max}$ = 0.5Å | 97% | 95% | 94% | 89% | 97% | 97% | 97% | 88% | 75% | 0.85 |
| 1:20, $D_{max}$ = 1Å | 93% | 88% | 85% | 85% | 92% | 93% | 93% | 76% | 57% | 0.86 |
| 1:50, $D_{max}$ = 0.1Å | 96% | 93% | 94% | 94% | 96% | 96% | 96% | 86% | 77% | 0.85 |
| 1:50, $D_{max}$ = 0.25Å | 97% | 91% | 95% | 94% | 96% | 96% | 96% | 90% | 81% | 0.85 |
| 1:50, $D_{max}$ = 0.5Å | 97% | 90% | 94% | 96% | 97% | 97% | 97% | 91% | 78% | 0.85 |
| 1:50, $D_{max}$ = 1Å | 89% | 81% | 84% | 81% | 89% | 89% | 89% | 73% | 54% | 0.85 |
| 1:100, $D_{max}$ = 0.1Å | 96% | 92% | 95% | 94% | 96% | 96% | 96% | 87% | 77% | 0.85 |
| 1:100, $D_{max}$ = 0.25Å | 96% | 94% | 95% | 93% | 96% | 96% | 96% | 84% | 77% | 0.85 |
| 1:100, $D_{max}$ = 0.5Å | 95% | 86% | 94% | 94% | 95% | 95% | 95% | 81% | 68% | 0.84 |
| 1:100, $D_{max}$ = 1Å | 62% | 72% | 42% | 31% | 56% | 62% | 62% | 46% | 8% | 0.84 |
| Performance with distortion factor conditioning | | | | | | | | | | |
| QM9 | 73% | 86% | 71% | 69% | 73% | 73% | 73% | 67% | 53% | 0.86 |
| GEOM | 10% | 20% | 40% | 80% | 80% | 100% | 100% | 60% | 0% | 0.70 |
| $GEOM_{no\ h}$ | 97% | 91% | 95% | 94% | 96% | 96% | 96% | 90% | 81% | 0.85 |
| ZINC | 90% | 96% | 83% | 87% | 87% | 90% | 90% | 75% | 63% | 0.84 |
| Performance with energy conditioning | | | | | | | | | | |
| QM9 | 57% | 40% | 56% | 57% | 56% | 57% | 57% | 53% | 19% | 0.78 |
| GEOM | 34% | 4% | 0% | 24% | 25% | 34% | 34% | 13% | 0% | 0.83 |
| $GEOM_{no\ h}$ | 98% | 96% | 96% | 96% | 98% | 98% | 98% | 88% | 84% | 0.85 |
| ZINC | 65% | 21% | 12% | 41% | 53% | 65% | 65% | 39% | 36% | 0.86 |
| GCDM | | | | | | | | | | |
| $GEOM_{no\ h}$ baseline | 100% | 92% | 100% | 100% | 95% | 100% | 100% | 90% | 83% | 0.89 |
| $GEOM_{no\ h}$ conditioned | 94% | 100% | 89% | 94% | 94% | 94% | 94% | 94% | 89% | 0.86 |
| ZINC baseline | 78% | 70% | 92% | 88% | 100% | 100% | 100% | 87% | 64% | 0.75 |
| ZINC conditioned | 100% | 100% | 100% | 100% | 100% | 100% | 100% | 78% | 78% | 0.76 |
| EquiFM | | | | | | | | | | |
| $GEOM_{no\ h}$ baseline | 97% | 93% | 96% | 95% | 100% | 100% | 100% | 91% | 57% | 0.85 |
| $GEOM_{no\ h}$ conditioned | 95% | 72% | 83% | 82% | 91% | 95% | 95% | 77% | 44% | 0.83 |
| ZINC baseline | 76% | 62% | 61% | 72% | 71% | 100% | 100% | 78% | 39% | 0.73 |
| ZINC conditioned | 96% | 97% | 94% | 95% | 96% | 96% | 96% | 90% | 87% | 0.85 |
